# Independent apical and basal mechanical systems determine cell and tissue shape in the *Drosophila* wing disc

**DOI:** 10.1101/2020.04.10.036152

**Authors:** Amarendra Badugu, Andres Käch

## Abstract

How cell shape and mechanics are organized in three dimensions during tissue morphogenesis is poorly understood. In the *Drosophila* wing imaginal disc, we examined the mechanical processes that determine the shape of epithelial cells. Since it has been known that basement membrane influences the mechanics intracellularly, we reexamined the material properties of the basement membrane with fluorescence and transmission electron microscopy in its native environment. Further, we investigated the effect on cell shape and tissue mechanics when disruptions were instigated at three different time scales: (1) short (seconds with laser cutting), (2) medium (minutes with drug treatments), and (3) long (days with RNAi interference). We found regions in which the basement membrane is much thicker and heterogeneous than previously reported. Disrupting the actin cytoskeleton through drug treatment affects cell shape only at the apical surface, while the shapes in the medial and basal surfaces were not altered. In contrast, when integrin function was inhibited via RNAi or basement membrane integrity was disrupted by drug treatment, the medial and basal cell shapes were affected. We propose that basement membrane thickness patterns determine the height and basal surface area of cells and the curvature of folds in the wing disc. Based on these findings and previous studies, we propose a model of how cell shapes and tissue properties were determined by highly local, modular apical and basal mechanics.

**Graphical abstract:** 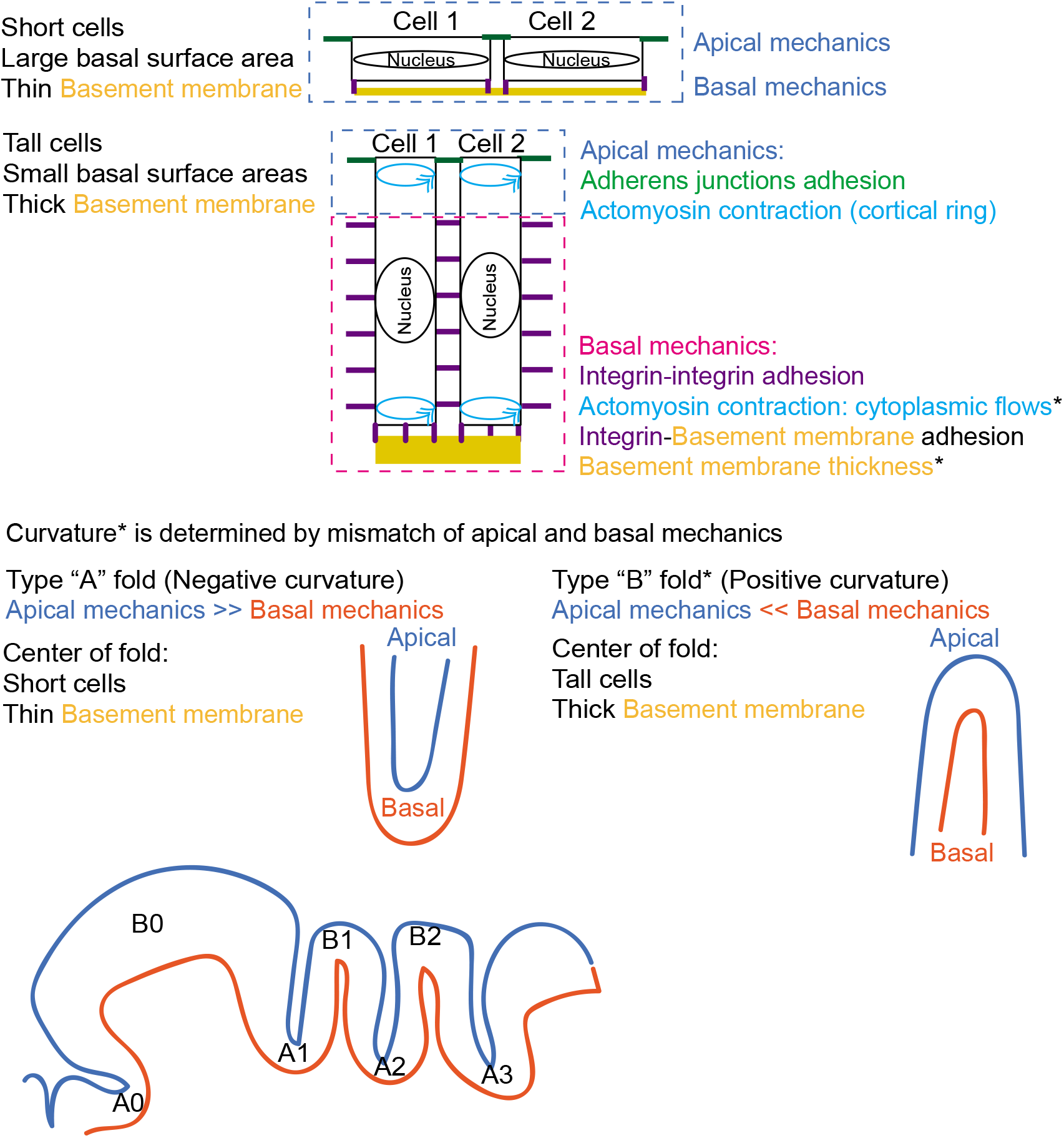

## Introduction

Our understanding of mechanical signals in tissue morphogenesis is still at its infancy compared to our understanding of biochemical signals (Campàs, 2016). Behind every movement in a cell or in a tissue there must be a physical force, even in the absence of instructive chemical signals (Campàs, 2016). Furthermore, the forces inside the cell operate at timescales of several orders magnitude smaller than those of biochemical signals. Advances in whole organ/deep tissue microscopy (Ahrens et al., 2013) (Helmchen and Denk, 2005) and three dimensional (3D) cell culturing techniques (Lee et al., 2007) were used to capture large image datasets of cell shapes. New image processing methods (Khan et al., 2014; Stegmaier et al., 2016) allow segmentation and quantification of cell shapes in 3D. Various studies have developed models to explain the shape of cells in 3D for simple epithelia, like those of the *Drosophila* embryo (Hannezo et al., 2014) or the *Drosophila* egg (Osterfield et al., 2013). Yet there is no logical biophysical framework explaining epithelial form in 3D for a complex epithelium such as the wing imaginal disc (M.C.Diaz de la Loza et al., 2017).

The extracellular matrix (ECM) is an organelle of the cell, which provides a barrier between the cell and the external microenvironment (Morrissey and Sherwood, 2015). The Basement Membrane (BM) is a special variant of ECM present in epithelial tissues and is composed of Laminin, Collagen IV, Perlecan, Nidogen and other proteins (Morrissey and Sherwood, 2015). In Drosophila, Collagen IV (Pastor-Pareja and Xu, 2011) and Perlecan (Zang et al., 2015) were synthesized in the fatbody and transported to the imaginal epithelia. In the Drosophila egg chambers, it has been shown that the Basement Membrane, through an anterior-posterior stiffness gradient, generates mechanical anisotropy that drives elongation of the tissue (Crest et al., 2017). The thickness of the ECM was assumed to be a uniformly thin sheet between 50 to 100 nm from ultrastructure studies conducted in mice (Kefalides et al., 1979, Halfter, W et al., 1987), whereas thicker BM was observed in the human vitreal surface of the retina, ranging from several nm during fetal stage to several μm at age 90 (Candiello et al., 2010). It was assumed that the laboratory animals like *Drosophila,* mice, rats, fish do not live long enough to accumulate thick BMs (Candiello et al., 2010). However detailed measurements of the basement membrane heterogeneity in its native state and models linking those patterns to tissue mechanics in an organ have not been made.

Molecularly, tissue mechanics can be divided into two systems, based on processes occurring apically and basally. Several studies have revealed the existence of mechanical events at the apical surface, here referred to as ‘apical mechanics’, which were defined by forces from cell-cell adhesion countered by an actin-based contractility (Farhadifar et al., 2007; Lecuit and Munro, 2011; Martin et al., 2009). Other studies have shown the existence of ‘basal mechanics’, playing an important role in determining cell shape through integrin-mediated contact with the basement membrane (Brabant et al., 1996; Domínguez-giménez et al., 2007) and via the composition of the Basement membrane (Pastor-Pareja and Xu, 2011; Zang et al., 2015). Although cell shape and mechanics at the apical surface have been quantitatively described at impressive detail, cell shape and mechanics at the basal surface was mostly ignored.

Therefore, we reexamined the properties of the basement membrane in its native state with electron microscopy and fluorescence imaging. The tissue and cell shape were explored in detail, taking into account both apical and basal surfaces. Further experiments were made to study how the mechanical processes at the apical and basal sides were coupled. We find that it is possible to experimentally sever the cell contact at the apical and basal surfaces separately in short time scales, and those respective effects were not transmitted through the entire cell. Further experiments with drug treatments and RNAi revealed similar effects were propagated in the medium and longer time scales. These results were integrated into a model of apical and basal mechanics.

## Results

### Morphology and cell shape in the wing disc

The larval wing disc (Fig 1A-A”’) is a three-dimensional bag-like structure that contains stratified columnar cells joined together by cuboidal cells and squamous cells known as peripodial cells and surrounded on the outside by a basement membrane (Fig 1B’, Fig S7D). The shapes of the cells were observed at the apical surface with the adherens junction protein bazooka (Fig 1B-B’, Fig S7A-A”, Fig S7B-C), the medial and basal surfaces with the marker resille (Fig S7A-A”, Fig S7B-C). The basal surface of the cells would be touching the inner side of the Basement membrane (Fig 1B’). Sectioning along the anterior posterior axis (green line representing the section in Fig 1A”’ and S7A”’) through the wing disc (Fig 1B-B’) reveal the curvature, which would be the measure of the value by which the surface deviates from a flat surface. The curvature can be categorized by the difference in the apical surface area (ASA) and the basal surface area (BSA). The categories are: Type A (TA), where ASA < BSA (Fig C); Type B (TB), where ASA > BSA (Fig C’); and Type C (TC) with ASA = BSA (Fig C”). The morphology of the wing disc in the anterior posterior axis was annotated with these categories (Fig 1C”’). The pouch was annotated as TB0/TC0 due to its structure comprising of both Type B (folding in the edges), and Type C (flat surface in the center) (Fig 1C”’, Fig S7B’, S7C’). The morphology in the dorsal ventral axis (Fig S7C, green dashed line in Fig S7A”’) was annotated with these categories (Fig S7C’).

**Fig 1.**
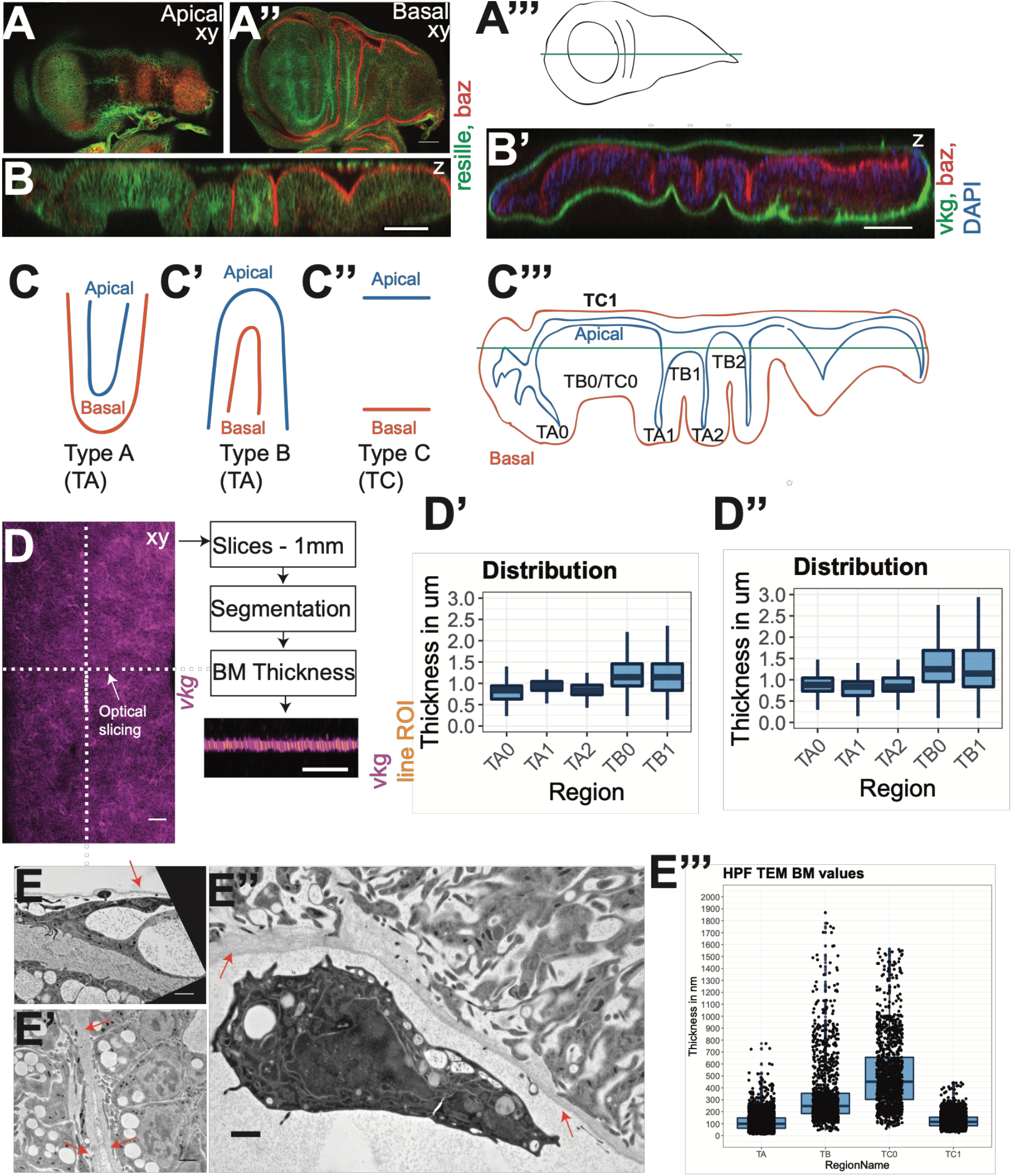
Apical, basal surfaces and patterns of basement membrane thickness patterns in the wing disc. (A-B) Resille-GFP (plasma membrane); baz-mcherry (apical surface). Representative images of wing disc (A) top down view of apical surface (A’) top down view of basal (B) lateral cross section along the anterior posterior axis. (A”) Graphical representation of the wing disc (top down). (B’) vkg-GFP (basal surface); baz-mcherry (apical surface); DAPI (nuclei). Cross section highlights the spatial separation between apical and basal surfaces. (C-C”) Curvature motifs (C) Type A, apical surface area < basal surface area (C’) Type B, apical surface area > basal surface area (C”) Type C, apical surface area = basal surface area. (C”’) Graphical representation of the wing disc (lateral cross section) with curvature motifs marked. Numbering starts with zero, left to right. (D-D”) Optical measurements of basement membrane (BM) thickness. (D) Method for automated optical sectioning and measurement of BM thickness from confocal images. Quantification of thickness patterns within a disc for BM. (D’) trol-GFP. (D’) vkg-GFP. (E-E”’) High pressure freezing TEM of basement membrane (BM) patterns. (E) BM on peripodial cells (red arrow). (E’) BM in TB fold (red arrows). (E”) BM in the pouch (red arrows). (E”’) Quantification of BM from HPF TEM. Scale bars, (A-B, B’, D): 50 μm; (E-E”’): 1 μm.

### Patterns of BM thickness in the wing disc

An image processing script was developed on confocal microscopy data (Fig 1D) that calculates the thickness profiles at different locations in the BM. To avoid observational errors due to the mismatch of the axial and lateral resolutions in confocal microscopy, this quantification method was applied only in regions that are parallel and closest to the cover slip. Different thickness profiles were observed (Fig 1D-D’).

For trol-GFP within a disc, the summary of BM thickness values for each of the regions were TA0 (0.837 (0.229) μm, mean (SD), N = 38960), TA1 (0.94 (0.325) μm, median (SD), N = 88970), TA2 (0.837 (0.221) μm, median (SD), N = 35135), TB1 (1.146 (1.421) μm, median (SD), N = 13796) and TB0/TC0 (Pouch) (1.25 (1.037) μm, median (SD), N = 215190) (Table S1).

For vkg-GFP within a disc, the BM thickness values for each of the regions were TA0 (0.881 (0.353) μm, median (SD), N = 99614), TA1 (0.837 (0.297) μm, median (SD), N = 89004), TA2 (0.837 (0.262) μm, median (SD), N = 63238), TB1 (1.767 (2.369) μm, median (SD), N = 4068) and TB0/TC0 (Pouch) (1.353 (2.083) μm, median (SD), N = 76275) (Table S2).

It should be noted that these measurements would be affected by the lateral resolution of the confocal microscope and subtraction of this resolution value (0.5 – 0.6 μm) from these measurements may thus give an approximate absolute value.

These thickness patterns were conserved between different wing discs for *trol-GFP* (Fig S1A-S1G’). and *vkg-GFP* (Fig S2A-S2G’). TB folds and TC0 (pouch) exhibits a thicker Basement Membrane than TA folds and TC1 (peripodial membrane).

### Confirmation of thickness patterns by super-resolution microscopy

Stimulated emission depletion (STED) microscopy is a super-resolution microscopy technique to bypass the diffraction limit for light microscopes (Hell and Wichmann, 1994). Leica TCS SP8 STED 3X applies STED in the axial direction providing 3D STED. Application of 3D STED microscopy on BM thickness patterns in the pouch and the peripodial membrane confirmed the above described thickness differences (Fig S4A-A”).

### Measurement of the thickness patterns with HPF and TEM

High-pressure freezing (HPF) combined with transmission electron microscopy (TEM) was often used to describe structures such as the Basement Membrane (BM) that was well preserved in its native condition. HPF and TEM was applied on the wing disc to determine the thickness of a well-preserved BM (Fig 1E-E”’). Thickness of BM in different sections of the wing disc was measured (Fig 1E, Fig S3A-B’). The summary statistics of the thicknesses are: TA folds (110.41 (66.76) nm, median (SD), N = 4730), TC1 (peripodial) (115 (53.39) nm, median (SD), N = 3990), TB folds (249 (286.33) nm, median (SD), N = 1333) and TB0/TC0 (pouch) (451 (297.59) nm, median (SD), N = 1305) (Table S3). BM thickness changed from TB fold to TA fold (Fig S3C-C’).

The BM of chemically fixed samples (Fig S3D) was measured for the pouch region (Fig S3D’) with a thickness of (92 (21.94) nm, median (SD), N = 810).

### Cell contact with basement membrane

Whereas Integrins mediate adhesion to the basement membrane, Talin regulates the strength of Integrin-adhesion to the cytoskeleton (Tanentzapf and Brown, 2006). Talin forms focal adhesion-like structures with a dot-like morphology at the basal surface in the wing disc (Tanentzapf and Brown, 2006) and behaves as a force vector sensor parallel to the surface of the membrane (Klapholz et al., 2015). Integrins formed fiber-like structures at the basal surface (Fig S5A-A’) whereas talin formed dot-like structures (Fig S5B-B’). By measuring the areas of the dot-like structures, we found that in the TC0 (pouch) cells, *talin* had smaller contact areas with the BM compared to the cells in the TA folds (Fig S5B”).

### Cell area at the basal surface

The basal cell cortices were segmented, and surface areas were calculated for TA1 (Fig 2A-A’) and TC0/TB0 (Pouch) (Fig 2B-B’, Fig SC). The basal cell areas were larger in the TA fold than in the TC0 pouch region. Distribution of basal surface areas indicates cells maintain a minimum area (Fig S8C’).

**Fig 2.**
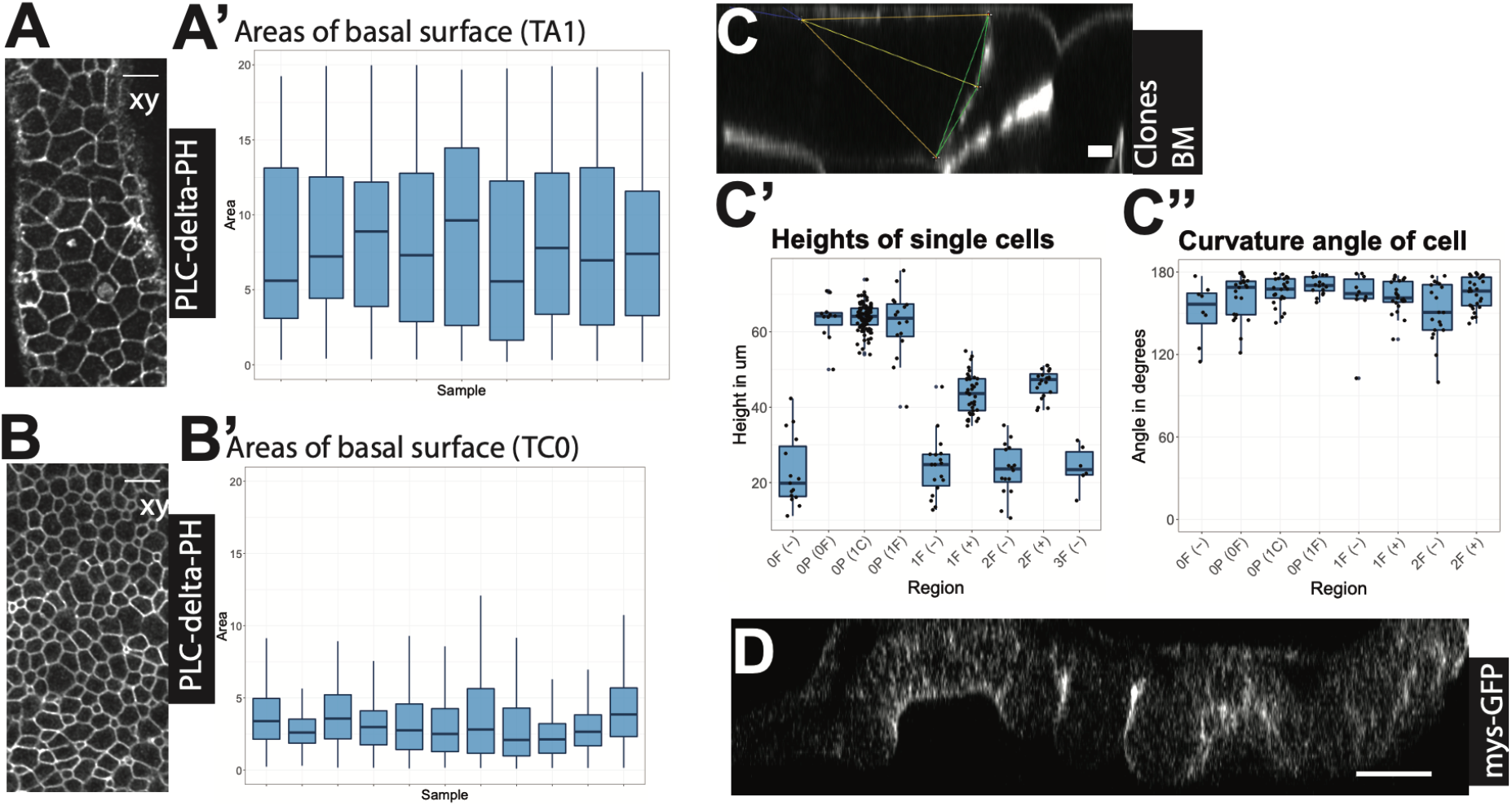
(A, B) *c765-*GAL4>UAS-PLCdelta-PH-GFP. Representative images of cell shapes at apical and basal surfaces. (A’, B’) Area distribution of segmented cell shapes. (C) *vkg-GFP* > *Act-Gal4;* UAS-GFP. Extraction of shape parameters from single cell clones in cross section. The point of intersection of yellow lines represent an angle of curvature. (C’) Quantification of heights in the wing disc from GFP tagged single cell clones. (C”) Quantification of angle of curvature in ‘the wing disc from GFP tagged single cell clones. (D) mys-GFP. Scale bars, (A, B, C): 50 μm; (E): 10 μm.

### Cell area at the apical and medial surfaces

The apical (Fig S8A) and medial (Fig S8B) cell cortices were segmented for the pouch region (TC0/TB0) and area distributions were calculated (Fig S8A’, Fig S8B’). This was only possible for a limited region (marked by rectangles in Fig S8A-B) due to the curvature of the pouch. The apical surface areas exhibit a normal distribution (Fig S8A’) whereas the medial surfaces exhibit a bimodal distribution (Fig S8B’).

### Shapes of single cells

If cells were columnar, they would be perfectly perpendicular to the basal and apical surfaces but single cell clones in the wing disc revealed that the cells were curved (Fig 2C). A cell under curvature would be taller than a cell that has zero curvature. By manually segmenting the heights of the curved cells, we found that cell heights were higher in the TC0 (pouch) and TB folds. Cell heights were lower in the peripodial membrane (TC1) and TA folds (Fig 2C). The curvature of single cells was found to be larger by (6.2 (2.8) %, median (SD), N = 30) for a cell height in the pouch and TB folds than zero curvature cells. Finally, Cells exhibited a wide distribution of angles and there was no discernible pattern (Fig 2C”).

### Integrin and actin levels

Integrin levels were higher in the TC0 (pouch) and TB folds. Integrin levels were lower in the peripodial membrane (TC1) and TA folds (Fig 2D).

Actin levels (Fig S5C) were higher at apical and basal surfaces in TC0 (pouch) (Fig S5C’) and TB1 (Fig S5C”). Actin levels were higher only at the apical surface and low at the basal surface in TA1 (Fig S5C”’).

### Disruption of basal and medial cell shape

It has previously been shown that Integrin-BM interactions were linked to the columnar-cuboidal transition (Domínguez-Giménez et al., 2007). We used polygonal packing as a metric of cell shape at different locations of cells (Fig S8A-C). If a tissue slice exhibits polygonal shapes that could be segmented, then we consider it ordered. To modify cell shape at longer time scales (48 hours), RNAi constructs were expressed in the dorsal compartment with the temperature sensitive UAS/Gal4 system. RNAi against *myospheroid (mys),* the aPS subunit of integrin showed a reduction in height for TC0 and TB1 (Fig 3A”’), except at the dorsal-ventral compartment boundary. RNAi against integrin has no effect in TA1. While the apical surface cell cortices (Fig 3A’) and some medial cortices (Fig 3A”) close to the apical surface were still ordered, the basal surface cell cortices were disordered (Fig 3A”’). In medium time periods (30 mins), the height of cells was reduced either by loss of BM with mild collagenase treatment (Fig 3C) or with trypsin treatment.

**Fig 3.**
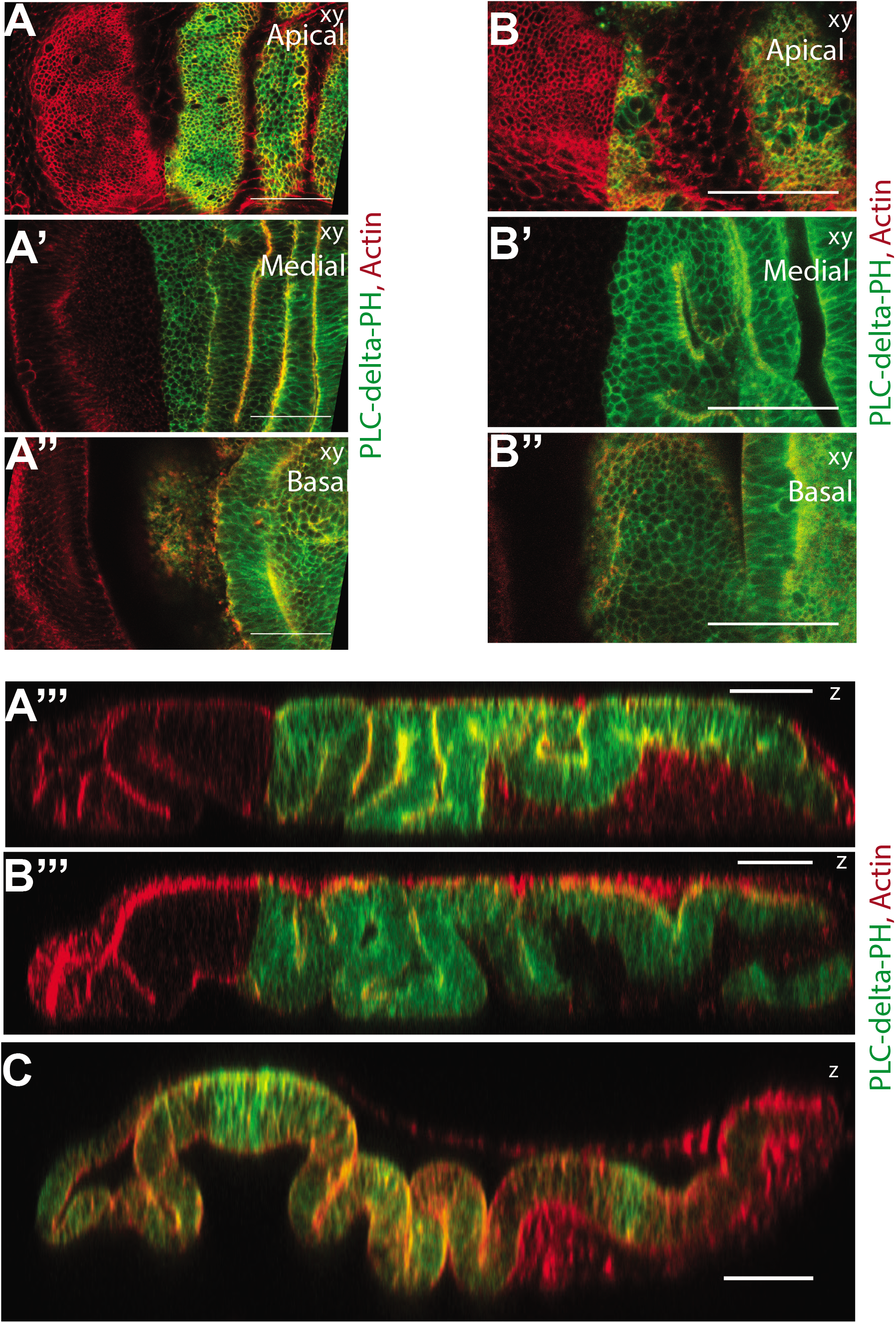
(A-A”’) *ap-*Gal4 tub-GAL80^ts^; UAS-PLCdelta-PH-GFP *> mys*^RNAi^ with actin staining. (B-B”’) *ap-*Gal4 tub-GAL80^ts^; UAS-PLCdelta-PH-GFP *> shg*^RNAi^ with actin staining. (A, A’) apical cell shapes. (A’, B’) medial cell shapes. (A”, B”) basal cell shapes. (A”’, B”’) cross sections. (C) Eversion with collagenase treatment for 30 min. Scale bars, (A-C): 50 μm

### Disruption of apical cell shape

RNAi against E-cadherin disrupted cell shape at the apical surface (Fig 3B’), while cell shape at the medial (Fig 3B’) and basal surfaces (Fig 3B”) remained ordered. E-cadherin caused a reduction of cell and tissue height (Fig 3B”’).

### Disruption of actin perturbs shape at apical surface and not at basal and medial surfaces

We treated wing discs with Latrunculin B the marine toxin that inhibits F-actin polymerization by forming complexes with actin monomers (Spector et al., 1989). Apical cell shape gets disrupted (Fig 4A-A’, 4B-B’). The treatment has no effect on cell shape in the medial (Fig 4C-C’) and basal surfaces (Fig 4D-D’, 4E), nor on the surface area of the *talin* patterning at the basal surface. Myosin accumulating in clumps at the apical surface (Fig S6A-A’) indicates disruption. The height of the cells in the pouch was found to be increased (Fig S6B-S6B”).

**Fig 4.**
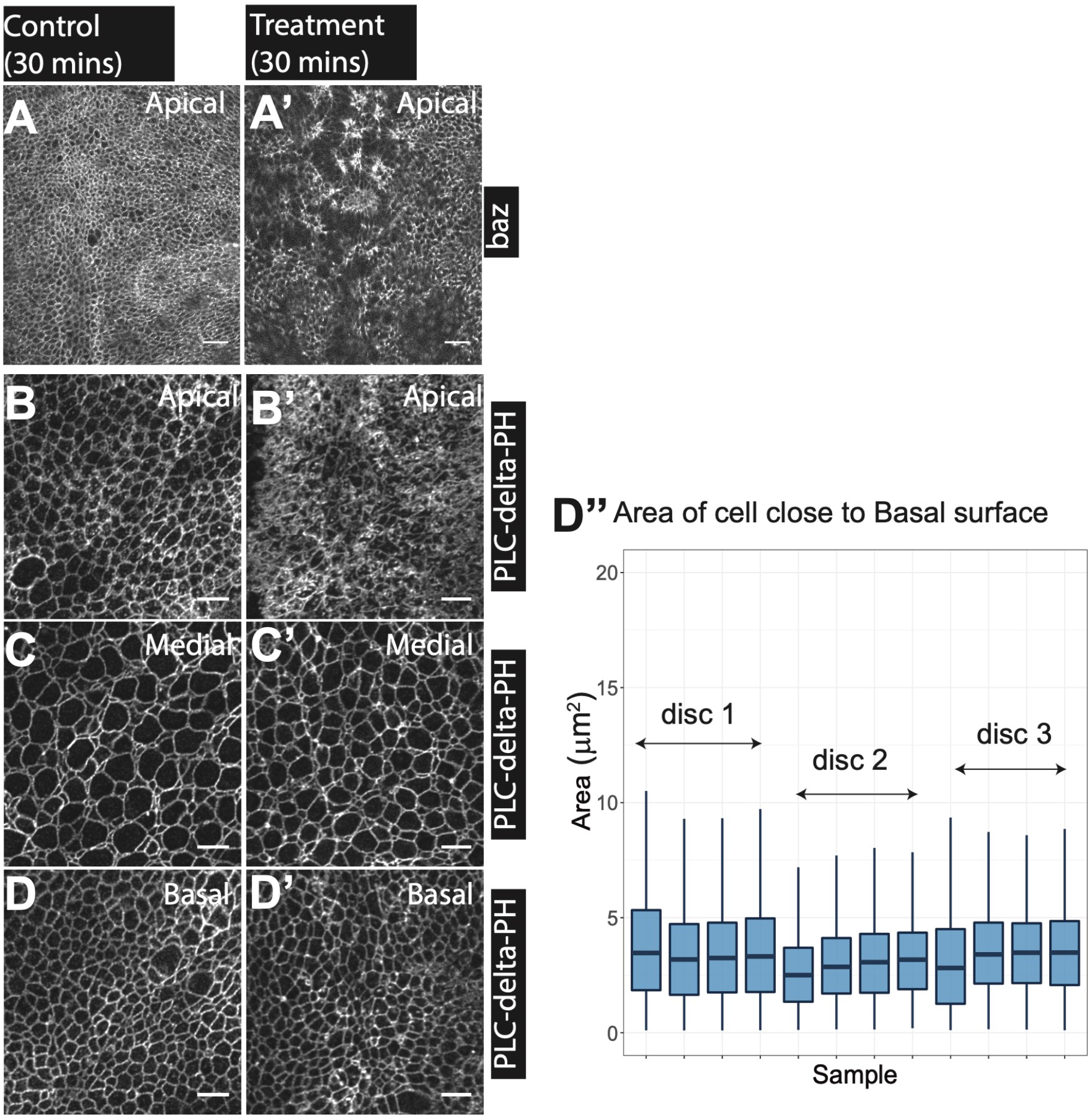
Effect of Latrunculin B (Lat B) on cell shape. (A-A’) baz-mcherry. Representative images of cell shapes at apical surface. (B-D’) *c765*-GAL4>UAS-PLCdelta-PH-GFP. Representative images of cell shapes at apical, medial, basal surfaces. (A, B) Apical cell shapes in control culture for 30 mins. (A’, B’) Apical cell shapes after treated with Lat B for 30 mins. (C) Medial cell shapes in control culture for 30 mins. (C’) Medial cell shapes after treated with Lat B for 30 mins. (D) Basal cell shapes in control culture for 30 mins. (D’) Basal cell shapes after treated with Lat B for 30 mins. (D”) Basal cell areas measured after Lat B treatment for 30 mins. Scale bars, (A-D’): 50 μm.

### Laser cutting of cell contact to apical and basal surfaces

UV laser ablation has previously been used to relax adherens junctions at the apical surface (Farhadifar et al., 2007). A two-photon laser was applied to ablate different contact surfaces of the cell to study the response in short time scales (seconds). We induced Act>Gal4 / UAS-GFP expressing clones. We ablated the cell contacts to the apical surface (formed by the contacts between the neighbors) (Fig 5A) and contact to the BM (Fig 5B). We observed an elastic response at the apical side of (5.72 (1.75) μm, median (SD), N = 13) and at the basal surfaces of (9.00 (1.72) μm, median (SD), N = 29), (Fig 5A’). This elastic response was followed by a rapid recovery very similar to a wound healing response.

**Fig 5.**
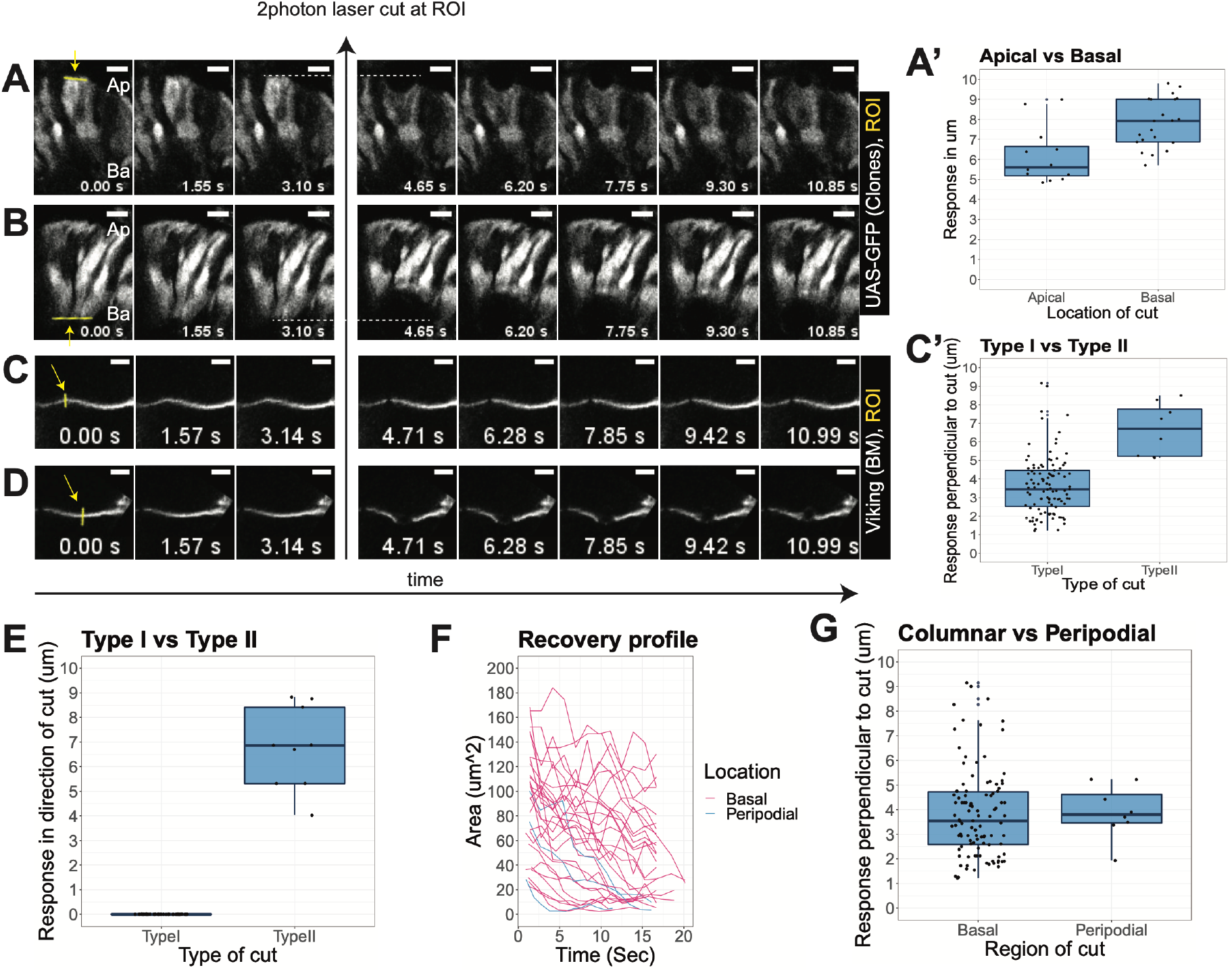
Response of 2-photon laser ablation on cell adhesion and basement membrane (BM). (A, B) *Act>Gal4;* UAS-GFP. (A-D) Yellow arrow in frame 1 is the region of interest (ROI) where the 2photon laser ablation is made between frames 3 and 4 indicated by the black arrow. (A) Response to ablation at apical contact of the cell. (B) Response to ablation at basal contact of the cell. (A’) Quantification of maximum retraction distance after ablation at apical and basal contacts respectively. (C, D) vkg-GFP. (C) Type I response to ablation at apical contact of the BM. (D) Type II response to ablation at apical contact of the BM. (C’) Quantification of maximum retraction distance of type I and type II responses in the axis perpendicular to the direction of the BM ablation. (E) Quantification of maximum retraction distance in the direction of the ablation of BM. (F) recovery profile over time for type II BM ablation. (G) Quantification of maximum retraction distance of peripodial and columnar cells perpendicular to the direction of the BM ablation. Scale bars, (A-D): 50 μm.

### Laser cutting of Basement membrane

A two photon laser was applied to ablate the BM (Fig 5C-D) to study the response in short time scales (seconds) on the pouch BM (3.56 (2.16) μm, median (SD), N = 108) and peripodial BM (4.15 (3.73) μm, median (SD), N = 10). We observed two types of responses: =Type I was a local relaxation after the cut perpendicular to the direction of the cut with no follow up of closing of the cut area (Fig 5C) and Type II was a relaxation in a circle followed by closing of the cut area (Fig 5D). Comparisons between Type I and Type II cuts were quantified (Fig 5C’, Fig 5E). The relaxation parallel to the direction of cut was (7.64 (2.27) μm, median (SD), N = 12) and relaxation perpendicular to the direction of cut was (7.9 (3.30) μm, median (SD), N = 12). Post cut, there was a recovery profile over time for type II BM ablation in peripodial and columnar BM (Fig 5F). Finally, there was a maximum retraction distance in peripodial and columnar BM ablation that was perpendicular to the direction of the ablation (Fig 5G).

## Discussion

### BM thickness patterns

In the *Drosophila* wing disc, we have observed BM thicknesses of several nm up to 2 μm – much larger than these previously reported measurements (Pastor-Pareja and Xu, 2011). Further, these thicknesses were not uniformly distributed in the wing disc; we have observed spatial patterning of the BM thickness in the disc: the TA folds attached to cuboidal cells and TC1 cells attached to peripodial cells have the lowest thickness values of BM. We were able to confirm that Measurement methods involving chemical dehydration causes a severe decrease in the thickness of the BM (Candiello et al., 2010).

### Cell-BM contacts, the link to cell shape and cytoskeleton

We have observed that both the TC0 (pouch) and TB folds attached to stratified columnar cells have the thickest BM. We also observed that cells in the TC0 and TB folds have higher heights, smaller basal contacts, smaller surface areas than TA folds and TC1 cells. Further, TC0 and TB folds had higher basal integrin intensity and higher basal actin levels. It has been shown through direct manipulation of BM components that BM mechanics are instructive to tissue morphogenesis and that the BM cannot be regarded as an inert scaffold that reflects the cell-intrinsic mechanics (Crest et al., 2017; Pastor-Pareja and Xu, 2011). Considering BM and Integrins were shown to be responsible for the cell height and columnar shape of the cells in the wing disc (Domínguez-giménez et al., 2007), it could be postulated that columnar cells through Integrins transform the thickness patterns into cell shape changes.

### Disruption of apical and basal cell shape

In short time scales (seconds), ablation of apical cell contacts and basal cell contacts causes an elastic response followed by a wound healing like response. In medium time scales (30 minutes), disruption of the BM causes reduction in height and an increase in surface area in TC0 and TB folds. Reduction of cell-cell adhesion via trypsin treatment causes a similar effect. Disruption of actin through Latrunculin B (Lat B) treatment causes loss of order at the apical surface, but the basal and medial cell shapes remain comparable to control samples. In longer time scales (48 hours), disruption of integrin expression causes a reduction of height and loss of tissue order at the basal surface in the pouch. Considering that both apical and medial surfaces have order, it was possible the apical mechanics was behaving as an independent entity. Similarly, loss of E-cadherin disrupts cell shape and loss of tissue order at the apical surface while basal and medial surfaces have cell shape and order.

### Modular apical and basal mechanics

Previous studies considered cells in a tissue as single mechanical entities and researchers have studied apical and basal mechanics molecularly in isolation of each other. The current models that consider both apical and basal mechanics were based on the assumption that they were related to each other by simple properties.

It had been reported that cells such as in gastric cancer that lose cadherins survive by upregulating the laminin/BM (Caldeira et al., 2015). When Integrins or BM were lost that causes a change in cell shape, cells can still survive, remain part of the epithelium and maintain adhesion with apical mechanics. Whereas the unicellular ancestor of metazoans contained all the components of cell adhesion (cadherins, integrins and extracellular matrix) (Niklas and Newman, 2016; Suga et al., 2013), the integrin adhesome was lost in both fungi and choanoflagellates (Karl J. Niklas and Newman, 2016; Sebé-pedrós et al., 2010), but both of them retain apical adhesion. The separation of basal and apical mechanics could not be identified separately if they were situated in tight proximity to each other, such as in cultured cells.

### Building a model of mechanics

Based on our findings and combined with previous knowledge, we propose a model of how tissue mechanics determine and maintain cell shape. Apical mechanics regulates and maintain the apical cell shape through adherens junctions and acto-myosin contraction of the cortical ring. Basal mechanics determine basal and medial cell shape through basement membrane (BM) thickness, strength of integrin-BM adhesion, acto-myosin contractions, and integrin-integrin adhesion.

Single cell traction force microscopy experiments reveal that the force/power and speed of contractility within the cell adapts to the external stiffness of the substrate (Fouchard et al., 2011). On stiff weakly deformable substrates, large contractile forces could be generated in the cell, whereas on soft easily deformable substrates contractile forces were low within the cell (Fouchard et al., 2011). Large contractile forces in regions with thicker BM could be responsible for generating smaller basal surface. Furthermore, apical contractility was shown to increase cell height through cytoplasmic flows (He et al., 2014). A similar mechanism could be proposed with basal contractility that increases heights of the cells in regions of larger BM thickness. Heights cannot be generated in regions with thinner BM with weaker contractile forces and consequently larger surface areas.

For a cell i, an apical unit (Fig 6A-6A’) could be defined as an entity which encompasses the contractile forces and adhesion from apical mechanics. Similarly, a basal unit (6B-6B’) could be defined as an entity which encompasses the contractile forces and adhesion from basal mechanics.

**Fig 6.**
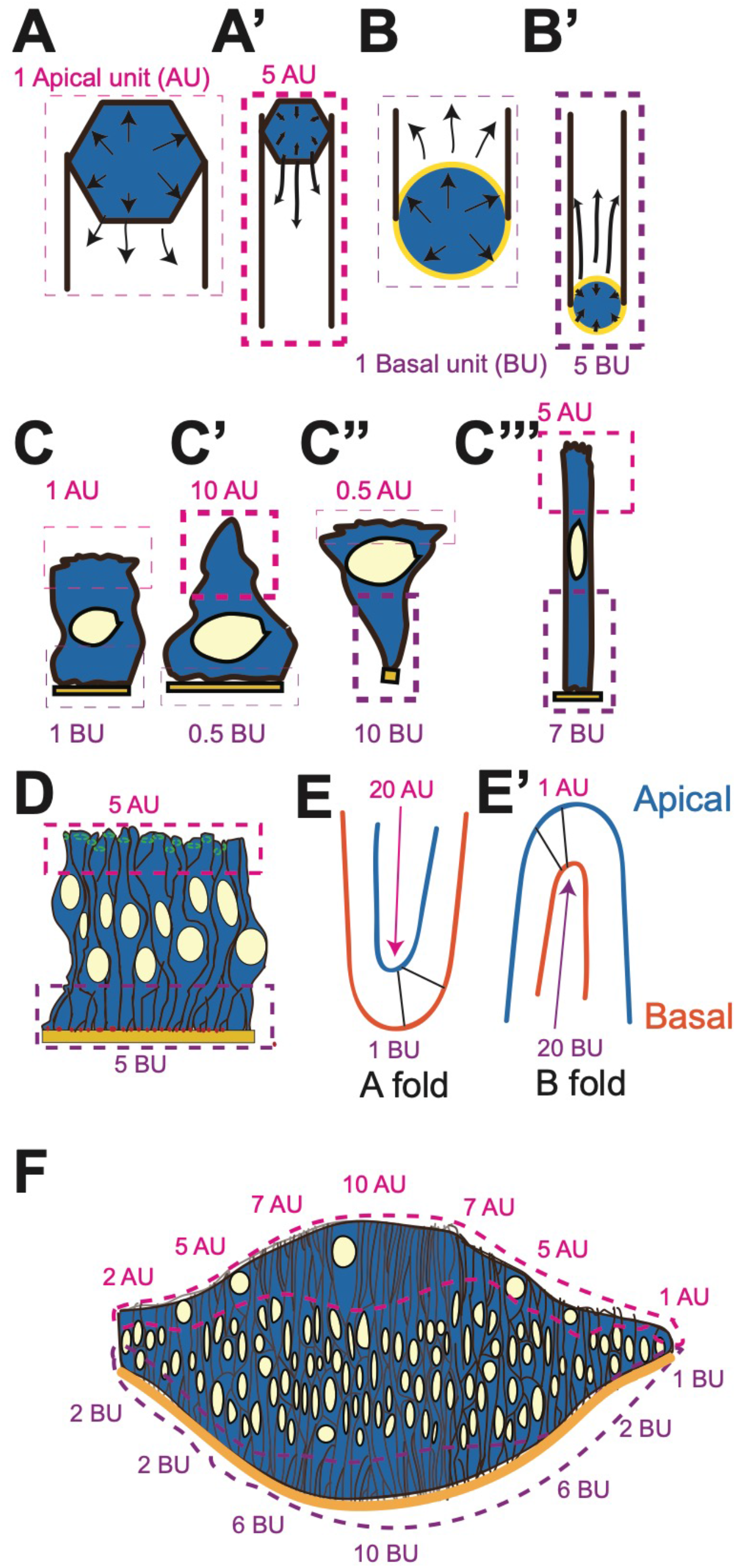
Model of cell and tissue shape with apical and basal units. (A-A’) Apical unit represents a hypothetical unit of the sum of the apical mechanics. (A) Low apical unit results in large apical surface area. (A’) High apical unit results in small apical surface area. (B-B’) Basal unit represents a hypothetical unit of the sum of the basal mechanics. (B) Low basal unit results in large basal surface area. (B’) High basal unit results in small basal surface area. (C-C”’) individual cell shapes were generated from combinations of differential apical and basal units. (D) If cells were modular subcomponents in a tissue, apical units and basal units in a tissue result in a flat surface. (E) TA fold is generated by a group of cells in the center of the fold having a very large apical unit compared to their basal unit. (E’) TB fold is generated by a group of cells in the center of the fold having a very large basal unit compared to their apical unit. (F) A smooth curvature is generated by a gradual reduction of apical and basal units.

The larger the unit will be, the smaller will be the cell area. The link between cell area and apical unit would follow the rule

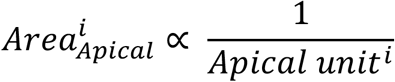

The link between cell area and basal unit would follow the rule

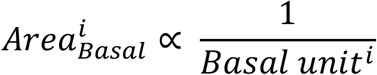

Modelling cells with differential apical and basal units predicts a variety of cell shapes (Fig 6C-C”’). If the values of apical and basal units were equal or coupled, a flat epithelial tissue would be generated (Fig 6D).

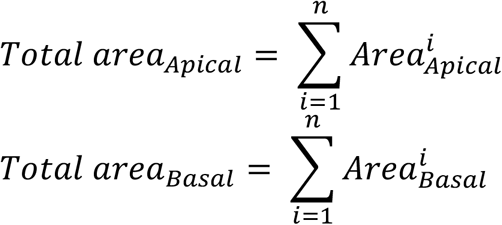

### Curvature

Curvature is the mismatch between the apical and basal surface areas.

For a TA fold,

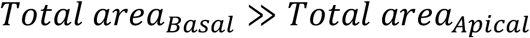

Centers of TA folds have a large apical unit compared to the basal unit (Fig 6E).

For a TB fold,

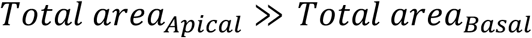

Centers of TB folds will have a large basal unit compared to the apical unit (Fig 6E’). We have observed that the centers of TB folds have thick BM and the thickness decreases as one moves towards the A fold. This thickness through acto-myosin contractility and integrin-based adhesion decreases the local surface area, generating a TB fold. Combining these two sets of observations, it was possible to predict tissue shapes with graded basal and apical units (Fig 6F).

In conclusion, we have identified thickness patterns of the Basement Membrane (BM) several μm larger than previously observed. We found that in regions of thick BM, cells had larger height, smaller basal surface area, smaller focal adhesion like kinase structure, higher levels of basal actin and Integrins compared to regions with thinner BM. By degrading the BM, we observed that cells in those regions change their shape to that of the cells in the thin BM regions. We propose that BM thickness is an instructive mechanical property that defines the local mechanics inside the cell. We observed that the difference in BM thickness correlates with the two types of curvature in the wing disc. We have manipulated mechanical behaviors and cell shapes at apical, medial and basal surfaces in the order of seconds, minutes and days through local laser ablations and drug and RNAi treatments of apical and basal mechanics. Finally, we propose a model of how cell shapes and tissue properties are determined with the apical and basal mechanics functioning as modular units.

## Methods

### Fly stocks and genetics

The following stocks were used: *Resille-GFP* (Martin et al., 2010, p. 200), *baz-mcherry* (McGill et al., 2009), vkg^G205^ (Morin et al., 2001), trol^G00022^ (Morin et al., 2001), *y[1] w[*]; L[1]/CyO; P{w[+mC]=UAS-PLCdelta-PH-EGFP}3/TM6B, Tb* (Bloomington 39693), *mys*^RNA1^ (Bloomington 33642)*, shg*^RNA1^ (VDRC 103962), *ap-*GAL4 tub-GAL80^ts^*/ CyO; Mkrs/ TM6B, Sp/ CyO;c-765-GAL4/ TM6B, mys-GFP* (Klapholz et al., 2015), *GFP-talin* (Klapholz et al., 2015). For UAS-GAL4 tub-GAL80^ts^ experiments, larvae were kept at 29c for 48 hours.

### Sample preparation

For living imaging, wing discs were mounted in an imaging chamber with a filter on top as described in (Zartman et al., 2013). For fixed imaging, samples were dissected in Ringers medium, were kept at 4% PFA overnight at 4°C before Immunostaining. Wing discs were mounted in VECTASHIELD (Vector Labs) inside a small cavity formed with reinforcement rings attached to the slide and avoiding compression from the cover slip.

### Immunofluorescence

1: 2000 SiR-Actin (Cytoskeleton, Inc).

### Drug treatments

20 mM stock solution of Latrunculin B (Lat B) in DMSO was diluted 1:400 in Schneider’s medium to a working concentration of 50 μM. Drug treatments were made for 30 minutes.

### Ultrastructure analysis of wing discs

#### High-pressure freezing

Wing discs have been dissected from L3 larvae in Schneider’s medium and quickly transferred onto carbon and polylysine coated 6 mm sapphire discs. For better orientation, a gold mask was added to the carbon-coated disc by sputtering prior to polylysine application. For imaging basal surfaces, the wing discs were oriented with the apical surface facing the sapphire disc. For imaging apical surfaces, the wing discs were oriented with the basal surface facing the sapphire discs. Extra medium was removed by using dental absorbent paper points. Sapphire discs containing the wing discs were transferred to the dedicated middle plate, covered with a 6 mm aluminum specimen carrier (150 μm cavity) wetted with 1-hexadecene and a 200 μm spacer ring, and immediately frozen using a Leica EM HPM100 high-pressure freezer (Leica Microsystems, Vienna, Austria) without using ethanol as a synchronizing medium.

Subsequently, samples were freeze-substituted in water-free acetone containing 1 % of osmium tetroxide for eight hours at −90°C, six hours at −60°C, four hours at −30°C and one hour at 0°C with transition gradients of 30°C per hour. Specimens were rinsed twice with water-free acetone, incubated in 1 % uranyl acetate in water-free acetone for one hour at 4°C, and rinsed again twice with water-free acetone, embedded in Epon/Araldite, 66 % in water-free acetone for two hours and 100 % for one hour prior to polymerizing at 60°C for 24 hours.

#### Chemical fixation

Wing discs were dissected from L3 larvae in Schneider’s medium and transferred onto 6 mm sapphire discs. Wing discs were fixed in 2.5 % glutaraldehyde for 6 hours at room temperature, rinsed with 0.1 M cacodylate buffer, incubated in 1 % osmium tetroxide in 0.1 M cacodylate buffer at room temperature followed by rinsing with pure water. Subsequently, samples were kept at 4°C for 1.5 hours in 1 % uranyl-acetate in water, followed by a water rinsing step and dehydration with a graded alcohol series (70 % for 20 minutes, 80 % for 20 minutes, 96 % for 45 minutes, 100 % for 15 minutes). Samples were then transferred to 100% propylene oxide for 30 minutes, incubated in 50% Epon/Araldite in propylene oxide at 4°C for two hours, and in 100% Epon/Araldite in a flat silicon embedding mold for one hour before polymerization at 60°C for 24 hours.

Ultra-thin sections were contrasted with Reynolds lead citrate and imaged in a CM 100 (at 80 kV) or Tecnai Spirit G2 (at 120 kV) transmission electron microscope (Thermo Fisher Scientific, Eindhoven, The Netherlands) using a side-mounted Gatan Orius 1000 digital camera and the Digital Micrograph software package (Gatan, Munich, Germany).

#### TEM images tile alignment

TrakEM2 (Cardona et al., 2012) was applied to align the TEM multiple tiles with least squares montage mode followed by elastic montage mode.

#### Confocal Imaging

Confocal images were acquired on a Zeiss LSM710. For fixed samples, confocal stacks were acquired with 40x 1.3 NA (oil) immersion objective with settings for deconvolution. For larger overviews, 25x (oil) objectives were used. For live samples, confocal stacks were acquired with 40x 1.2 NA water immersion objective. For imaging basal side, discs were mounted basal side closer to the objective.

#### 2 photon image acquisition

2 photon imaging were acquired on an Olympus Fluoview 1000 MPE.

#### Image analysis

Images were deconvolved with Huygens Professional version 17.04 (Scientific Volume Imaging, The Netherlands, http://svi.nl).

The confocal stacks were sliced in top and left directions and then the individual slices were thresholded and segmented; a custom script in Fiji (Schindelin et al., 2012) was developed that calculates the thickness following the shape of the BM from slices. The bending angles of the cells were calculated (Fig 2D) by applying method from (Feduccia, 1993).

#### Cell shape segmentation and feature extraction

A median filter of 0.5 was applied on the images with the cell cortices. “Find Maxima” plugin in Fiji was applied to segment cell outlines. “Analyze Particles” plugin in Fiji was applied to extract features from the segmented outlines.

## Author contributions

AB conceived the project, performed the experiments and data analysis and wrote the manuscript. AK performed the ultrastructure preparation and TEM imaging.

## Acknowledgments

I would like to thank Konrad Basler for the position in his laboratory and support as my PhD advisor. I would like to thank Simon Tanaka, Robert Witte and Dominik Eder for their valuable inputs, feedback and discussion during the writing of this manuscript. I would like to thank Bloomington Drosophila Stock Center, Vienna Drosophila RNAi Center, Stefan Luschnig and Nick Brown for reagents. I would like to thank Carmen Kaiser for TEM preparation. I would like to thank Center for Microscopy and Image Analysis, University of Zurich for their TEM and 2-photon. This work was supported by the University of Zurich.

## Bibliography

Ahrens, M.B., Orger, M.B., Robson, D.N., Li, J.M., Keller, P.J., 2013. Whole-brain functional imaging at cellular resolution using light-sheet microscopy 10. https://doi.org/10.1038/NMETH.2434

Brabant, M.C., Fristrom, D., Bunch, T.A., Brower, D.L., 1996. Distinct spatial and temporal functions for PS integrins during Drosophila wing morphogenesis 3317, 3307–3317.

Caldeira, J., Figueiredo, J., Brás-pereira, C., Carneiro, P., Moreira, A.M., Pinto, M.T., Relvas, J.B., Carneiro, F., Barbosa, M., Casares, F., Janody, F., Seruca, R., 2015. E-cadherin-defective gastric cancer cells depend on Laminin to survive and invade 24, 5891–5900. https://doi.org/10.1093/hmg/ddv312

Campàs, O., 2016. A toolbox to explore the mechanics of living embryonic tissues. Semin. Cell Dev. Biol. https://doi.org/10.1016/j.semcdb.2016.03.011

Cardona, A., Saalfeld, S., Schindelin, J., Arganda-Carreras, I., Preibisch, S., Longair, M., Tomancak, P., Hartenstein, V., Douglas, R.J., 2012. TrakEM2 Software for Neural Circuit Reconstruction. PLoS ONE 7, e38011. https://doi.org/10.1371/journal.pone.0038011

Crest, J., Diz-Muñoz, A., Chen, D.-Y., Fletcher, D.A., Bilder, D., 2017. Organ sculpting by patterned extracellular matrix stiffness. eLife 6. https://doi.org/10.7554/eLife.24958

Domínguez-giménez, P., Brown, N.H., Martín-bermudo, M.D., 2007. Integrin-ECM interactions regulate the changes in cell shape driving the morphogenesis of the Drosophila wing epithelium. https://doi.org/10.1242/jcs.03404

Farhadifar, R., Röper, J.-C., Aigouy, B., Eaton, S., Jülicher, F., Ro, J., Aigouy, B., Eaton, S., 2007. Article The Influence of Cell Mechanics, Cell-Cell Interactions, and Proliferation on Epithelial Packing. Curr. Biol. CB 17, 2095–2104. https://doi.org/10.1016/j.cub.2007.11.049

Feduccia, A., 1993. Evidence from Claw Geometry Indicating Arboreal Habits of Archaeopteryx Science 259, 790 LP–793.

Fouchard, J., Mitrossilis, D., Asnacios, A., 2011. Acto-myosin based response to stiffness and rigidity sensing. Cell Adhes. Migr. 5, 16–19. https://doi.org/10.4161/cam.5.1.13281

Hannezo, E., Prost, J., Joanny, J., 2014. Theory of epithelial sheet morphology in three dimensions 111, 27–32. https://doi.org/10.1073/pnas.1312076111

He, B., Doubrovinski, K., Polyakov, O., Wieschaus, E., 2014. Apical constriction drives tissue-scale hydrodynamic flow to mediate cell elongation. Nature 508, 392–396. https://doi.org/10.1038/nature13070

Hell, S.W., Wichmann, J., 1994. Breaking the diffraction resolution limit by stimulated emission: stimulated-emission-depletion fluorescence microscopy. Opt. Lett. 19, 780–782. https://doi.org/10.1364/OL.19.000780

Helmchen, F., Denk, W., 2005. Deep tissue two-photon microscopy 2. https://doi.org/10.1038/NMETH818

Karl J. Niklas, Newman, S.A., 2016. Multicellularity: Origins and Evolution. MIT Press.

Khan, Z., Wang, Y., Wieschaus, E.F., Kaschube, M., 2014. Quantitative 4D analyses of epithelial folding during Drosophila gastrulation 2895–2900. https://doi.org/10.1242/dev.107730

Klapholz, B., Herbert, S.L., Wellmann, J., Johnson, R., Parsons, M., Brown, N.H., 2015. Alternative Mechanisms for Talin to Mediate Integrin Function. Curr. Biol. 25, 847–857. https://doi.org/10.1016/j.cub.2015.01.043

Lecuit, T., Munro, E., 2011. Force Generation, Transmission, and Integration during Cell and Tissue Morphogenesis. https://doi.org/10.1146/annurev-cellbio-100109-104027

Lee, G.Y., Kenny, P.A., Lee, E.H., Bissell, M.J., 2007. Three-dimensional culture models of normal and malignant breast epithelial cells 4, 359–365. https://doi.org/10.1038/NMETH1015

Martin, A.C., Gelbart, M., Fernandez-Gonzalez, R., Kaschube, M., Wieschaus, E.F., 2010. Integration of contractile forces during tissue invagination. J. Cell Biol. 188, 735–749. https://doi.org/10.1083/jcb.200910099

Martin, A.C., Kaschube, M., Wieschaus, E.F., 2009. Pulsed contractions of an actin-myosin network drive apical constriction. Nature 457, 495–9. https://doi.org/10.1038/nature07522

McGill, M.A., McKinley, R.F.A., Harris, T.J.C., 2009. Independent cadherin–catenin and Bazooka clusters interact to assemble adherens junctions. J. Cell Biol. 185, 787–796. https://doi.org/10.1083/jcb.200812146

Morin, X., Daneman, R., Zavortink, M., Chia, W., 2001. A protein trap strategy to detect GFP-tagged proteins expressed from their endogenous loci in Drosophila. Proc. Natl. Acad. Sci. 98, 15050–15055. https://doi.org/10.1073/pnas.261408198

Morrissey, M.A., Sherwood, D.R., 2015. An active role for basement membrane assembly and modification in tissue sculpting. J. Cell Sci. 128, 1661–1668. https://doi.org/10.1242/jcs.168021

Osterfield, M., Du, X., Schu, T., Wieschaus, E., Shvartsman, S.Y., 2013. Article in the Developing Drosophila Egg 400–410. https://doi.org/10.1016/j.devcel.2013.01.017

Pastor-Pareja, J.C., Xu, T., 2011. Shaping Cells and Organs in Drosophila by Opposing Roles of Fat Body-Secreted Collagen IV and Perlecan. Dev. Cell 21, 245–256. https://doi.org/10.1016/j.devcel.2011.06.026

Poodry Clifton A., S.H.A., 1971. Intercellular Adhesivity and Pupal Morphogenesis in Drosophila melanogaster 9, 1–9.

Schindelin, J., Arganda-Carreras, I., Frise, E., Kaynig, V., Longair, M., Pietzsch, T., Preibisch, S., Rueden, C., Saalfeld, S., Schmid, B., Tinevez, J.-Y., White, D.J., Hartenstein, V., Eliceiri, K., Tomancak, P., Cardona, A., 2012. Fiji: an open-source platform for biological-image analysis. Nat. Methods 9, 676–682. https://doi.org/10.1038/nmeth.2019

Sebé-pedrós, A., Roger, A.J., Lang, F.B., King, N., Ruiz-trillo, I., 2010. Ancient origin of the integrin-mediated adhesion and signaling machinery. https://doi.org/10.1073/pnas.1002257107/-/DCSupplemental.www.pnas.org/cgi/doi/10.1073/pnas.1002257107

Stegmaier, J., Amat, F., Lemon, W.C., Teodoro, G., Mikut, R., Keller, P.J., Stegmaier, J., Amat, F., Lemon, W.C., Mcdole, K., Wan, Y., Teodoro, G., Mikut, R., 2016. Real-Time Three-Dimensional Cell Segmentation in Large-Scale Microscopy Data of Developing Technology Real-Time Three-Dimensional Cell Segmentation in Large-Scale Microscopy Data of Developing Embryos. Dev. Cell 36, 225–240. https://doi.org/10.1016/j.devcel.2015.12.028

Suga, H., Brown, M.W., Kramer, E., Chen, Z., Mendoza, A. De, Sebe, A., Carr, M., Kerner, P., Vervoort, M., Manning, G., Lang, B.F., Russ, C., Haas, B.J., Roger, A.J., Nusbaum, C., 2013. The Capsaspora genome reveals a complex unicellular prehistory of animals 1–9. https://doi.org/10.1038/ncomms3325

Tanentzapf, G., Brown, N.H., 2006. An interaction between integrin and the talin FERM domain mediates integrin activation but not linkage to the cytoskeleton. Nat. Cell Biol. 8, 601–606. https://doi.org/10.1038/ncb1411

Zang, Y., Wan, M., Liu, M., Ke, H., Ma, S., Liu, L.-P., Ni, J.-Q., Carlos Pastor-Pareja, J., 2015. Plasma membrane overgrowth causes fibrotic collagen accumulation and immune activation in Drosophila adipocytes. eLife 4. https://doi.org/10.7554/eLife.07187

Zartman, J., Restrepo, S., Basler, K., 2013. A high-throughput template for optimizing *Drosophila* organ culture with response-surface methods. Development 140, 2848–2848. https://doi.org/10.1242/dev.098921

