## Supplementary Data for "Independent apical and basal mechanical systems determine cell and tissue shape in the *Drosophila* wing disc"

### Fig S1

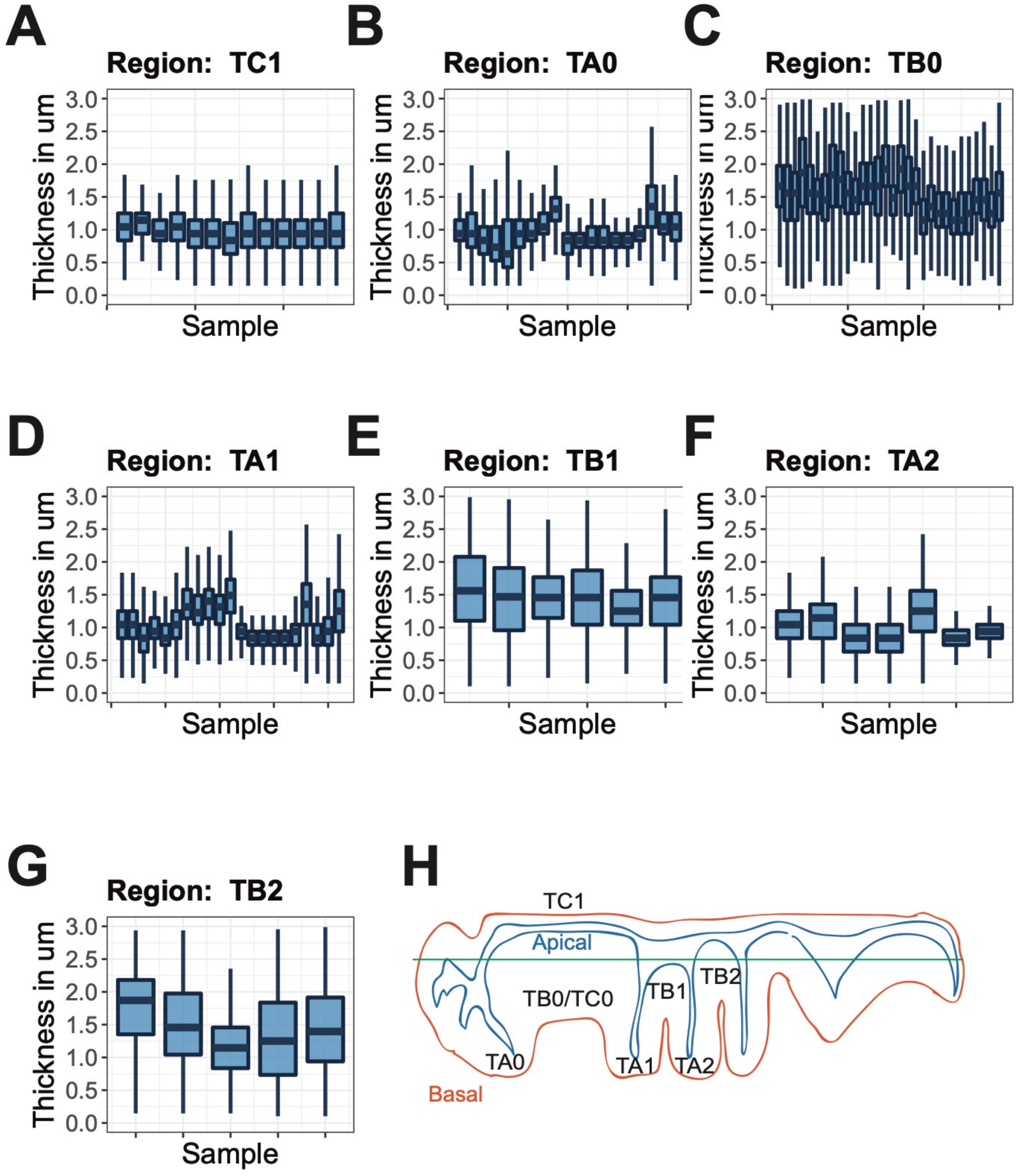


Fig S1. Boxplots of the distribution of BM thickness measurements for different regions in the wing disc from confocal datasets of Perlecan (trol-GFP). Each Box represents a different wing disc. (A) TC1 (Peripodial). (B) TA0 fold. (C) TB0/TC0 (Pouch). (D) TA1 fold. (E) TB1 fold. (F) TA2 fold. (G) TB2 fold.

### Fig S2

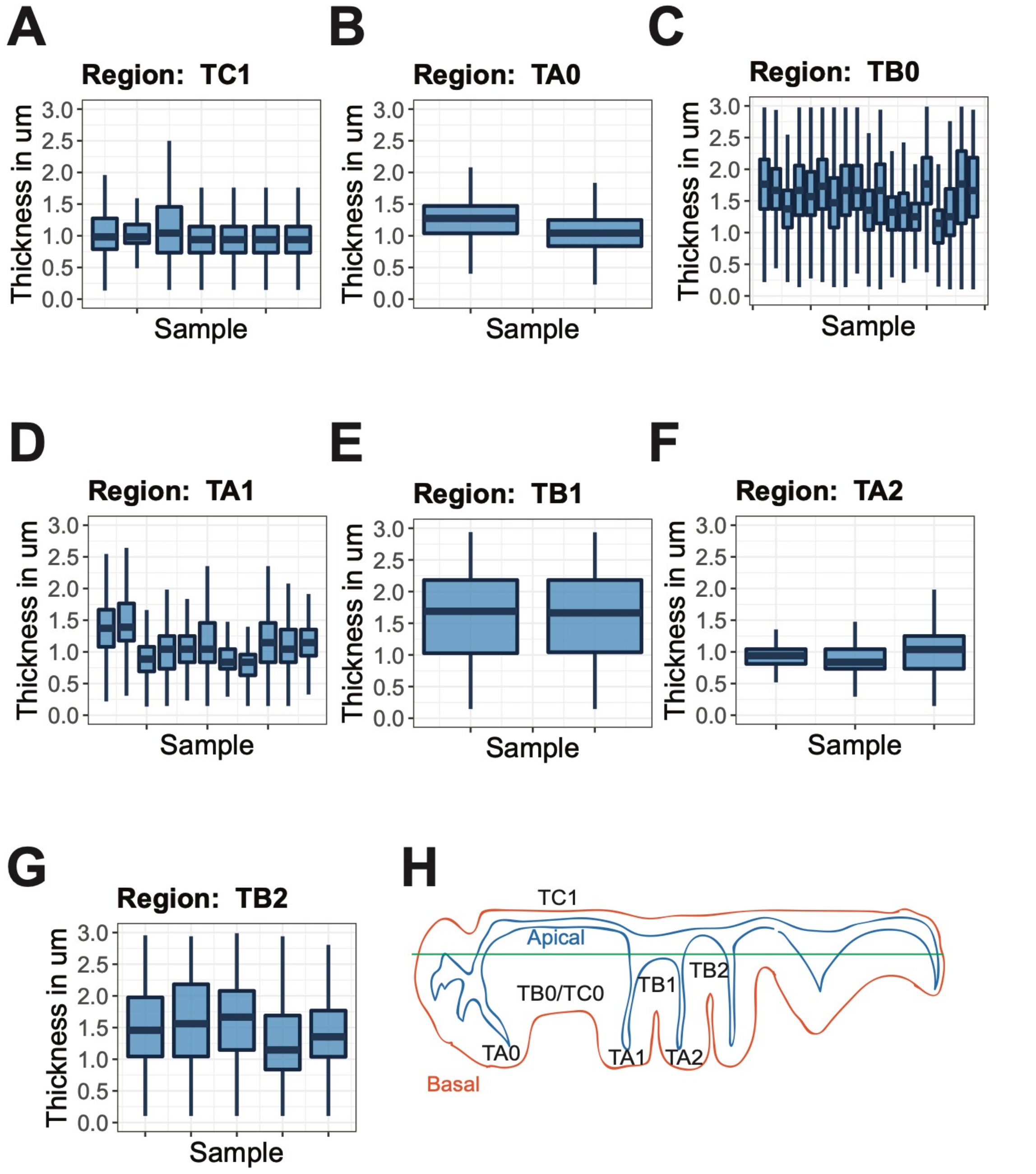


Fig S2. Boxplots of the distribution of BM thickness measurements for different regions in the wing disc from confocal datasets of Collagen IV (vkg-GFP). Each Box represents a different wing disc. (A) TC1 (Peripodial). (B) TA0 fold. (C) TB0/TC0 (Pouch). (D) TA1 fold. (E) TB1 fold. (F) TA2 fold. (G) TB2 fold.

### Fig S3

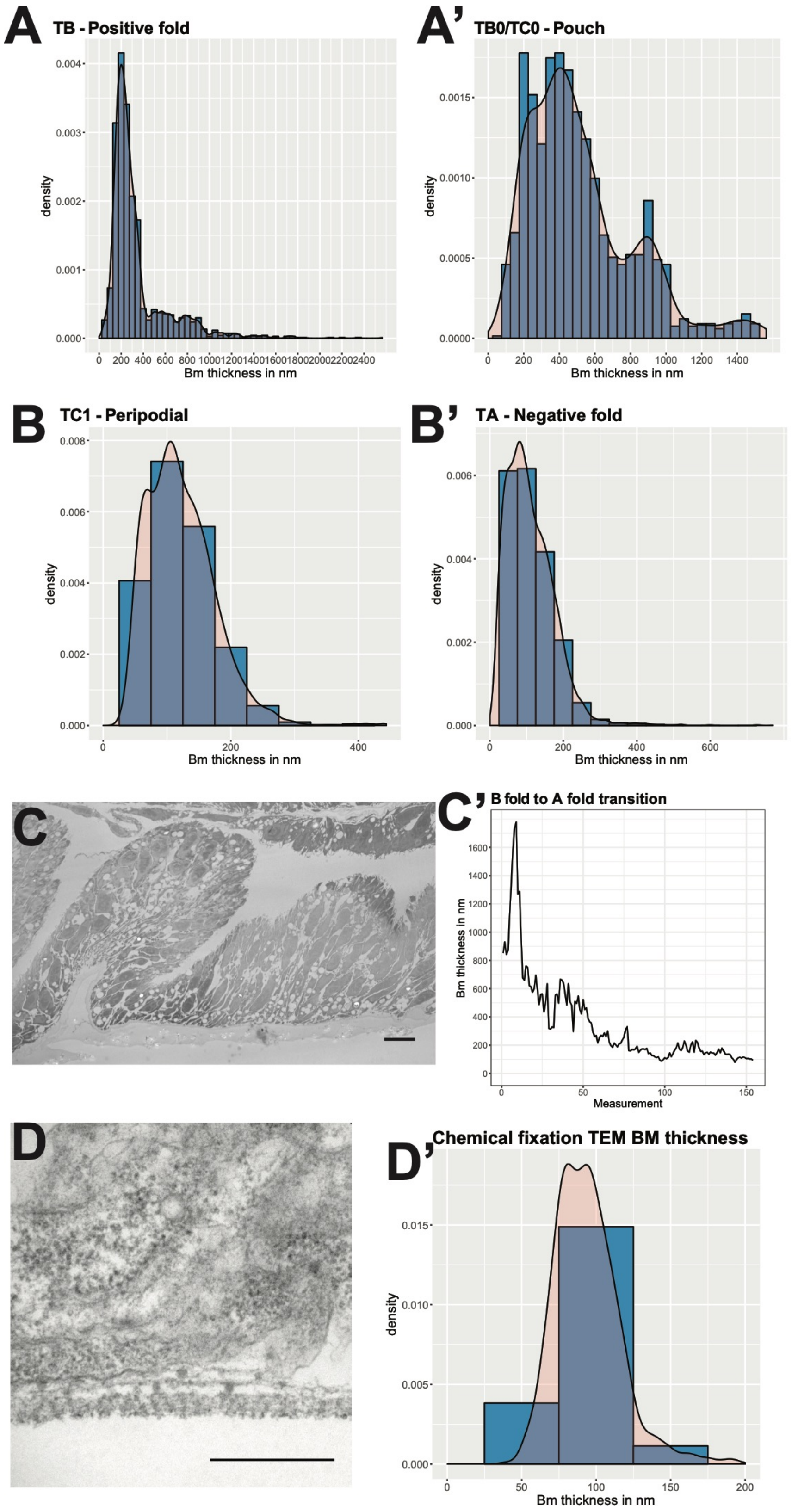
A
TB - Positive fold
density
Bm thickness in nm
A'
TB0/TC0 - Pouch
density
Bm thickness in nm
B
TC1 - Peripodial
density
Bm thickness in nm
B'
TA - Negative fold
density
Bm thickness in nm
C
C'
B fold to A fold transition
Bm thickness in nm
Measurement
D
D'
Chemical fixation TEM BM thickness
density
Bm thickness in nm

Fig S3. Quantification of basement membrane (BM) thickness patterns from High Pressure Freezing Transmission Electron Microscopy (HPF-TEM) and Chemically Fixed Transmission electron microscopy (CF-TEM) images. (A-B') Histograms of measured BM thickness from HPF-TEM images. (C) HPF-TEM image of transition between TB fold and TA fold. (C') Quantification of Change in thickness of BM from TB fold to TA fold in HPF-TEM. (D) Example CF-TEM image from the pouch region. (D) Quantification of the thickness from different regions of CF-TEM images. Scale bars, (C): 5 μm; (D): 0.5 μm.

### Fig S4

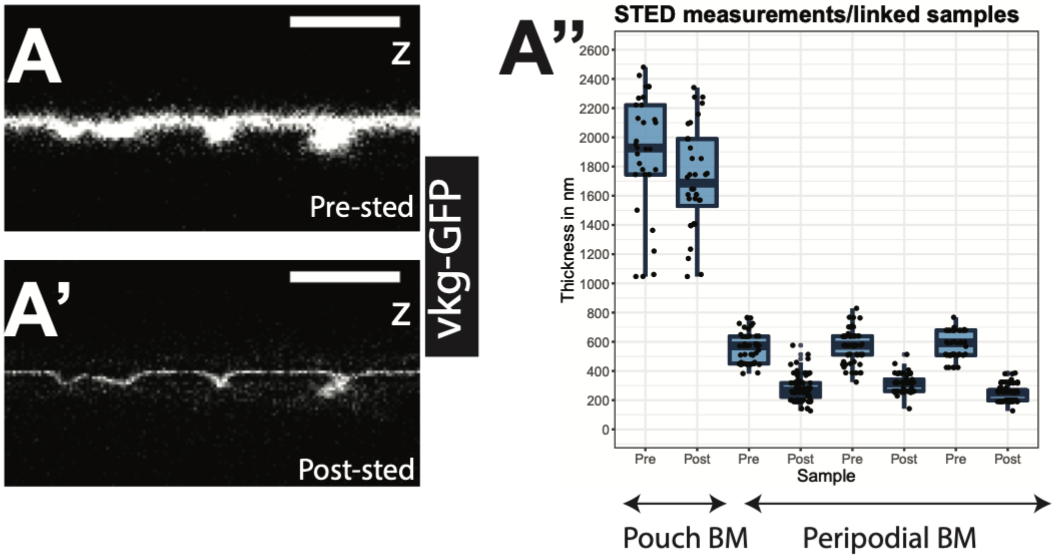


Fig S4. Quantification of basement membrane (BM) thickness patterns with STED microscopy. (A) Representative cross section image of vkg-GFP (basement membrane) pre 3D STED. (A') Representative cross image of vkg-GFP (basement membrane) after 3D STED (post STED). (A'') quantification of thickness pre and post STED. Scale bars, (A-A'): 5 µm.

Fig S5

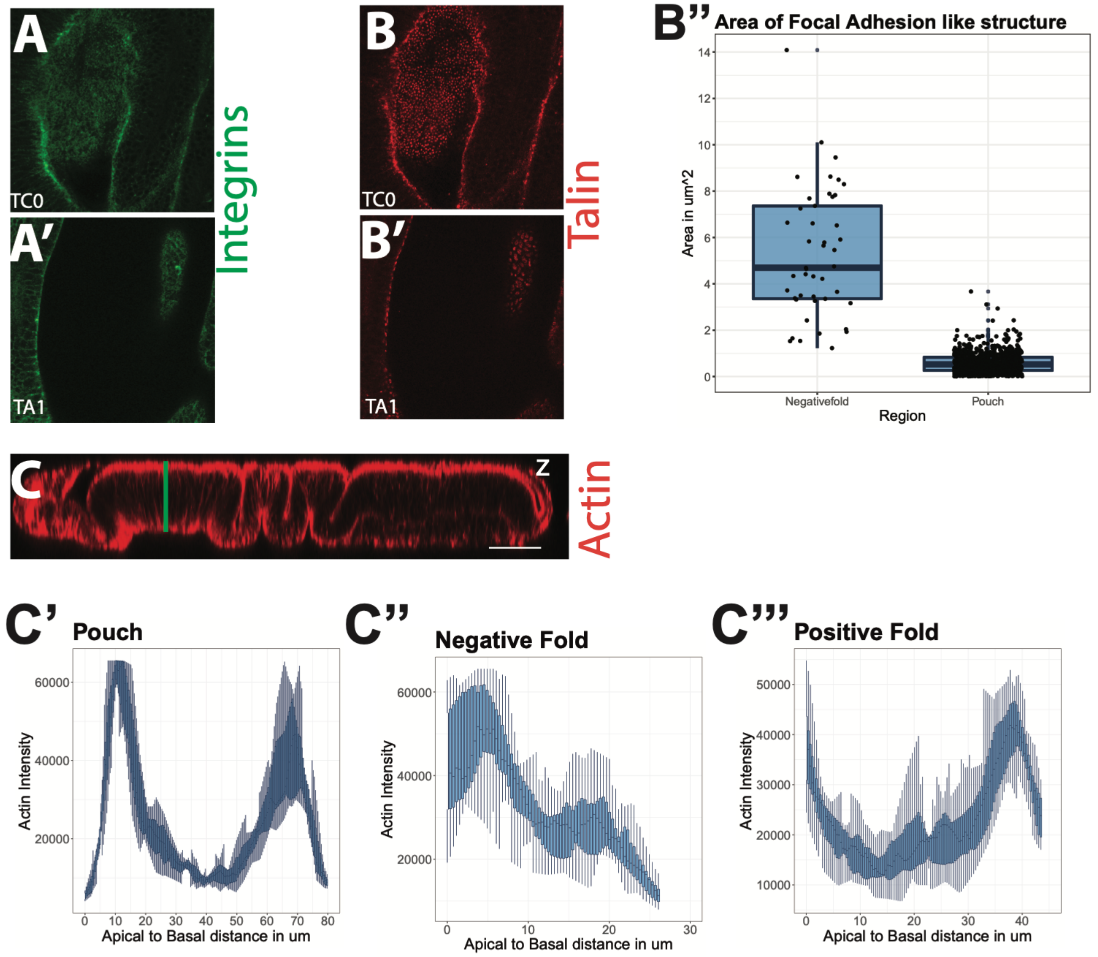


Fig S5. Integrins, talin and actin organization in wing disc. (A) mys-GFP. (B) talin-mcherry forms Focal adhesion kinase (FAK) like structures. (B) Quantification of areas of FAK like structures in the pouch and negative fold. (D) Actin staining in a representative cross section. (C'-C''') Quantification of actin signal from apical to basal surface. (C) Pouch region. (C') Type A fold with negative curvature. (C'') Type B fold with positive curvature. Scale bars, (A-D): 50 µm.

### Fig S6

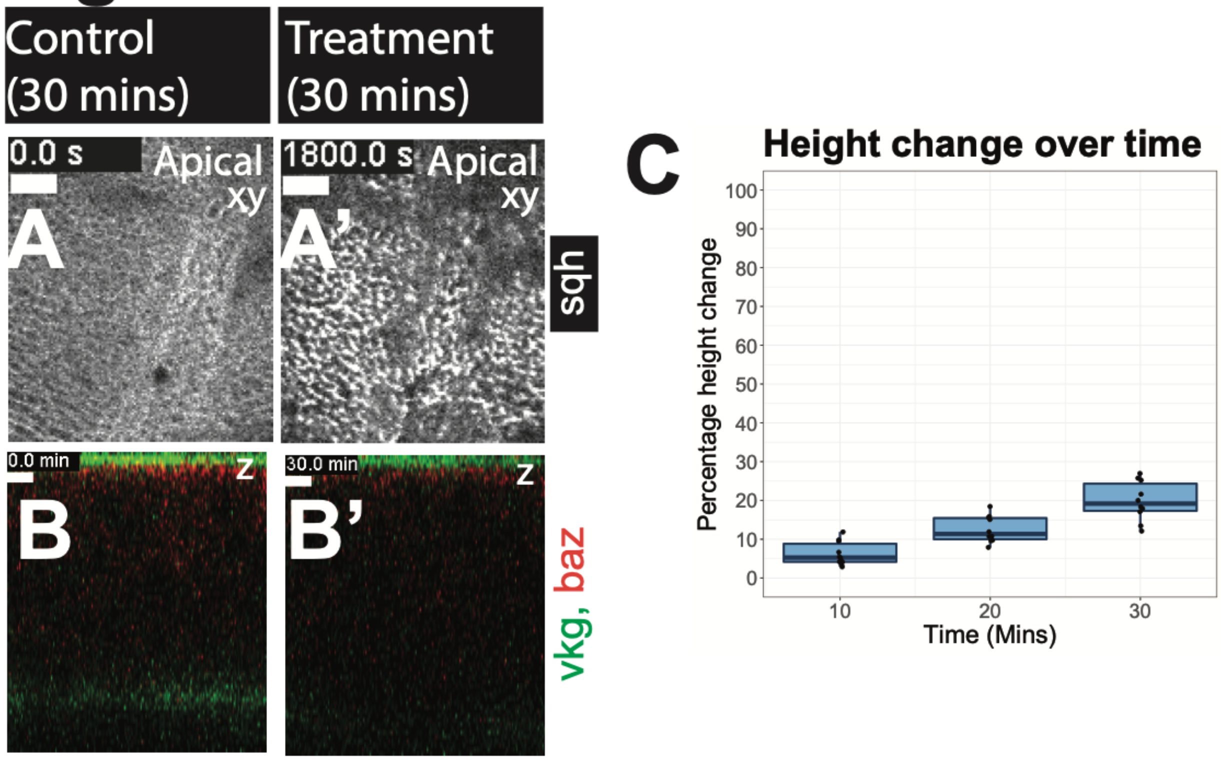


Fig S6. Effect of Latrunculin B (Lat B) on myosin and cell height. (A-A') sqh-GFP. (A) Myosin in culture for 30 mins (control). (A') myosin after treated with Lat B for 30 mins (treatment). (B-B') vkg-*GFP*; baz-mcherry; (B) representative image of cross section before LatB. (B') cross section after treated with Lat B for 30 mins. (B'') Quantification of measurements of cell height (between basement membrane and apical surface) change over time with Lat B treatment. Scale bars, (A-B'): 50 µm.

### Fig S7

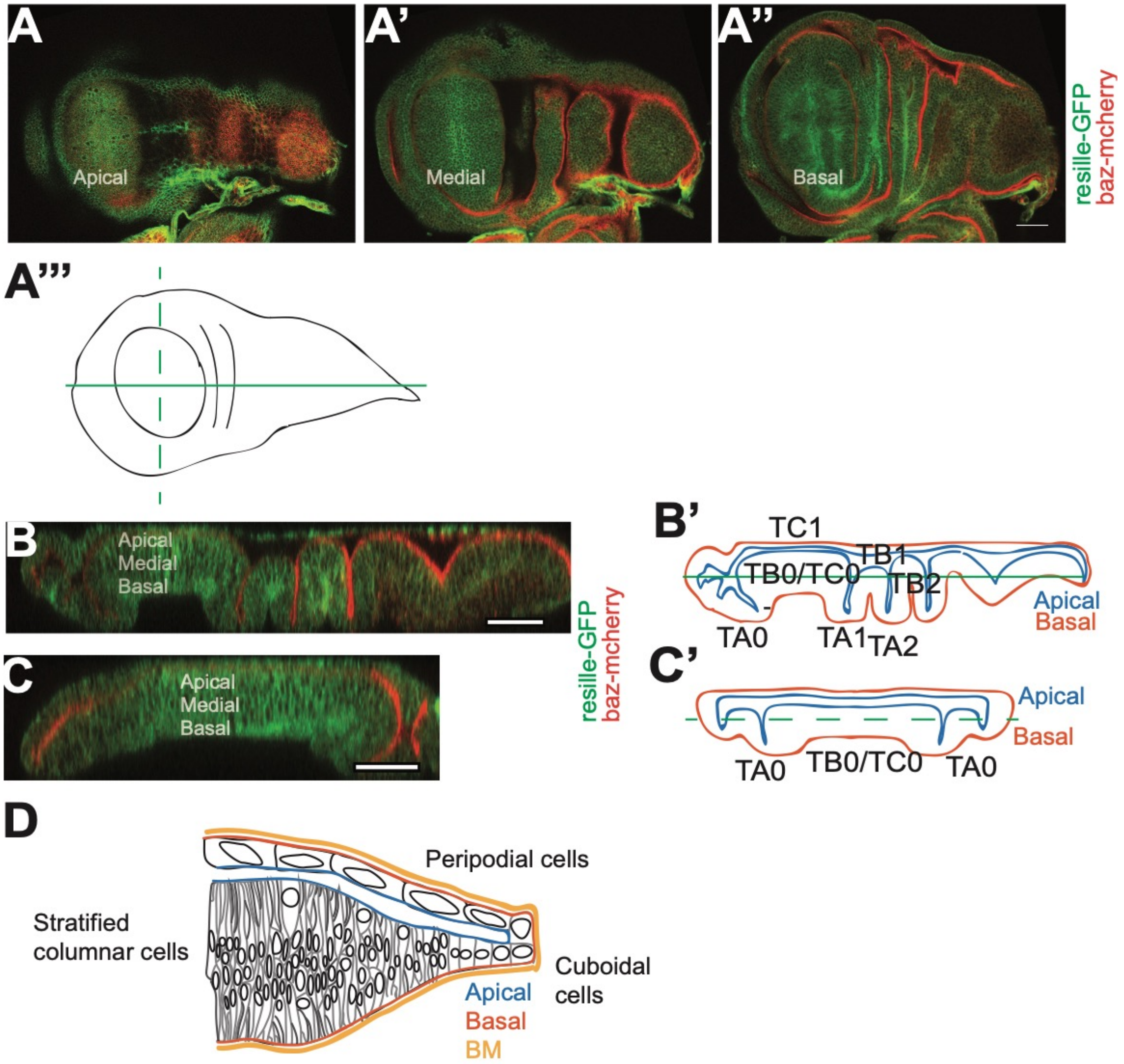


Fig S7. Morphologies and cell types in the wing disc. (A-C) resille-GFP (plasma membrane); baz-mcherry (apical surface). Representative images of the labelled section. (A) top down view of apical section. (A') top down view of medial section. (A'') top down view of basal section. (A''') graphical representation of the wing disc (top down), solid green line represents section through anterior posterior axis and dashed green line represents section through dorsal ventral axis. (B-B') lateral cross section in the anterior posterior axis. (C-C') lateral cross section in the dorsal ventral axis. (B', C') graphical representations of the sections with labelled curvature motifs: Type A (apical surface area < basal surface area), Type B (apical surface area > basal surface area), and Type C (apical surface area = basal surface area). (D-D'') Graphical representation of cell types: stratified columnar cells, cuboidal cells, and peripodial cells surrounded by Basement Membrane (BM). Scale bars, (A-A'', B, C): 50 µm.

### Fig S8

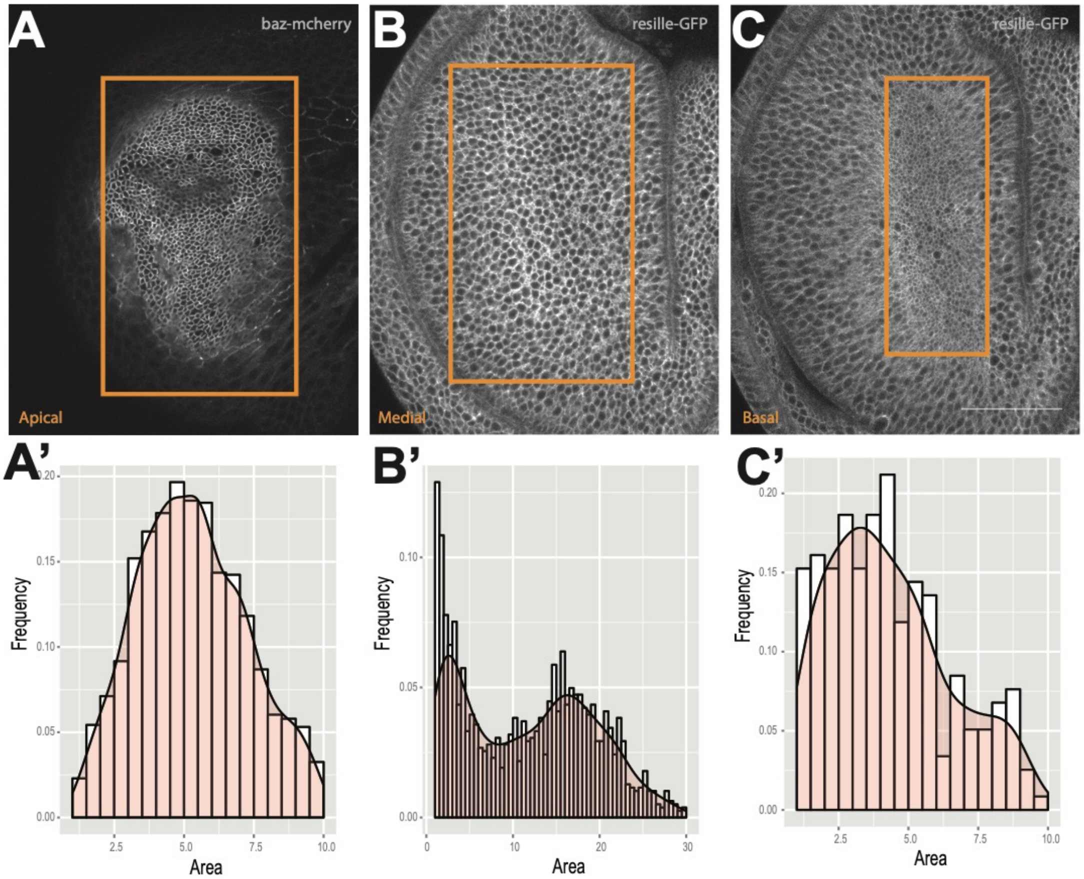


Fig S8. Cell shapes in top down sections in the pouch region. (A-C) Representative images of top-down sections. (A) baz-mcherry (apical surface). (B-C) resille-GFP (plasma membrane). (A) top down view of apical section. (A') top down view of medial section. (A'') top down view of basal section. (A'-C') Quantification of segmented cell areas of (A-C) respectively. Scale bars, (A-A''): 50 µm.

| Region | Median Thickness (µm) | Standard Deviation (µm) | Measurement Numbers |
|---|---|---|---|
| TA0 | 0.837 | 0.229 | 38960 |
| TA1 | 0.94 | 0.325 | 88970 |
| TA2 | 0.837 | 0.221 | 35135 |
| TB1 | 1.146 | 1.421 | 13796 |
| TB0/TC0 | 1.25 | 1.037 | 215190 |

Table S1. Optical thickness measurements of BM within a representative disc: trol-GFP.

| Region | Median Thickness (µm) | Standard Deviation (µm) | Measurement Numbers |
|---|---|---|---|
| TA0 | 0.881 | 0.353 | 99614 |
| TA1 | 0.837 | 0.297 | 89004 |
| TA2 | 0.837 | 0.262 | 63238 |
| TB1 | 1.767 | 2.369 | 4068 |
| TB0/TC0 | 1.353 | 2.083 | 76275 |

Table S2. Optical thickness measurements of BM within a representative disc: vkg-GFP.

| Region | Median Thickness (nm) | Standard Deviation (nm) | Measurement Number |
|---|---|---|---|
| TA folds | 110.41 | 66.76 | 4730 |
| TC1 | 115 | 53.39 | 3990 |
| TB folds | 249 | 286.33 | 1333 |
| TB0/TC0 | 451 | 297.59 | 1305 |

Table S3. Thickness measurements of BM in High pressure frozen with transmission electron microscopy samples.
